# EMO: Predicting Non-coding Mutation-induced Up- and Down-regulation of Risk Gene Expression using Deep Learning

**DOI:** 10.1101/2023.11.21.568175

**Authors:** Zhe Liu, Yihang Bao, Weichen Song, Guan Ning Lin

**Affiliations:** Shanghai Mental Health Center, Shanghai Jiao Tong University School of Medicine, School of Biomedical Engineering, Shanghai Jiao Tong University, Shanghai, China; Shanghai Key Laboratory of Psychotic Disorders, Shanghai, China; Bio-X Institutes, Key Laboratory for the Genetics of Developmental and Neuropsychiatric Disorders (Ministry of Education), Collaborative Innovation Center for Brain Science, Shanghai Jiao Tong University, Shanghai, China

## Abstract

The challenge of understanding how alterations in non-coding DNA regulate gene expression is substantial, with far-reaching consequences for the advancement of human genetics and disease research. Accurately predicting the up- and down-regulation of gene expression quantitative trait loci (eQTLs) offers a potential avenue to accelerate the identification of associations between non-coding variants and phenotypic traits. However, current methods for predicting the impact of non-coding mutations on gene expression changes fail to predict the sign of eQTLs accurately. Additionally, the requirement for tissue-specific training models within these methods restricts their applicability, especially when extending predictive abilities to single-cell resolution. In this study, we present EMO, an innovative transformer-based pre-trained method, designed to predict the up- and down-regulation of gene expression caused by single non-coding mutations using DNA sequences and ATAC-seq data. EMO extends the effective prediction range up to 1Mbp between the non-coding mutation and the transcription start site (TSS) of the target gene. It demonstrates competitive prediction performance across various variant TSS distances and surpasses the state-of-the-art structure. To assess its effectiveness, EMO was fine-tuned using eQTLs from two brain tissues for external validation. We also evaluated EMO’s transferability to single-cell resolution by fine-tuning it on eQTLs from six types of immune cells, achieving satisfactory results in each cell type (AUC > 0.860). Furthermore, EMO displayed promising potential in analyzing disease-associated eQTLs.

## 1 Introduction

Deciphering gene expression regulation and the effects of genome variation is crucial in human genetics and precision medicine[1-3]. The previous studies on predicting the deleteriousness of genetic variation within individual genomes have primarily concentrated on protein-altering variants[4, 5]. However, non-coding variants constitute most human genetic variation[6, 7]. The vast noncoding mutation space presents a significant challenge for deciphering risks, particularly in understanding how noncoding DNA impacts gene expression in different cell types[8]. By studying how mutations in non-coding regions affect gene expression, we can better understand the regulatory networks of genes[9]. Also, non-coding mutations may also affect epigenetic marks such as DNA methylation and histone modification[10], which in turn affect gene expression. Understanding these effects can help reveal complex epigenetic regulatory mechanisms.

Several computational methods have been developed to predict the effect of non-coding mutations on gene expression, such as DeepSea[11], Basenji2[12], Expecto[1], and Enformer[8]. However, these methods are limited to predicting gene expression changes in regulatory regions within 100kb upstream or downstream of the mutation, even though the range of cis-regulation can extend up to 1Mb[13]. Moreover, it has been reported that current genomic deep learning models inadequately explain personal transcriptome variation[14, 15]. These methods train models for specific tissues, highlighting a lack of generalizability. They can only make predictions for cell types and tissues in their training data and are not transferable to new cell types or tissues. Prior studies have also indicated that these models often inaccurately predict the direction of cis-regulatory genetic variation’s impact on expression[8, 14].

Certain non-coding mutations may result in either the overexpression or insufficient expression of pathogenic genes[16, 17]. Analyzing or predicting the influence of these non-coding mutations on the expression patterns of disease-causing genes can facilitate the understanding of the genetic basis of diseases and contribute to revealing their molecular mechanisms. Thus, predicting the up- and down-regulation of gene expression quantitative trait loci (eQTL) could accelerate the identification of associations between non-coding variants and phenotypic traits. However, even the most advanced model, Enformer, has struggled with accurately predicting the sign of eQTLs, particularly for regulatory ranges beyond 1kbp[8].

In this study, we introduced EMO, a pre-trained model designed to predict the direction of effect of the single cis-regulatory non-coding mutations on gene expression[14] using DNA sequences and ATAC-seq data, which can be fine-tuned to new tissues and cell types without training specific models. It is a cost-effective, segmented model capable of making predictions within a regulatory range of 1Mb, offering interpretability.

We evaluated EMO’s prediction performance in tissue-specific fine-tuning against Enformer, finding that the representations obtained from pre-training can enhance the accuracy and stability of tissue-specific eQTL sign prediction tasks. Additionally, we visualized the embeddings generated by EMO through representation learning and conducted an ablation study to assess the independent contributions of each model component. To demonstrate EMO’s fine-tuning and generalization capabilities to new tissues, we externally validated it using eQTLs from two new brain tissues not included in the training set. It was observed that the pre-trained EMO’s embeddings enhanced the rationality of prediction outcomes compared to end-to-end predictions. Moreover, through transfer learning, we tested EMO on single-cell eQTLs across six different cell types, achieving results in all (AUC > 0.860). Notably, EMO also accurately predicted two disease-associated eQTLs, underscoring its potential in unraveling the mechanisms of disease-risk SNPs.

## 2 Results

### 2.1 EMO structure and performance in short- and long-range prediction

Although some methods are available to annotate the impact of DNA mutations on gene expression, there is still room for improvement in predicting the direction of this impact[14]. As a complement to existing methods, EMO was developed as a pre-trained architecture that receives DNA sequences and ATAC-seq as inputs to output a trend in the effect of DNA mutations on gene expression (categorized as either ‘up-regulation’ or ‘down-regulation’, Figure 1, “Methods”), thereby further annotating the impact of mutations. Specifically, DNA sequences are encoded in a One-Hot format. The DNA sequences from the variant transcription start site (TSS) to the variant and the surrounding 51 base pairs (bp) of DNA are introduced into the model, together with the corresponding ATAC-seq values at the corresponding location.

**Figure 1.**
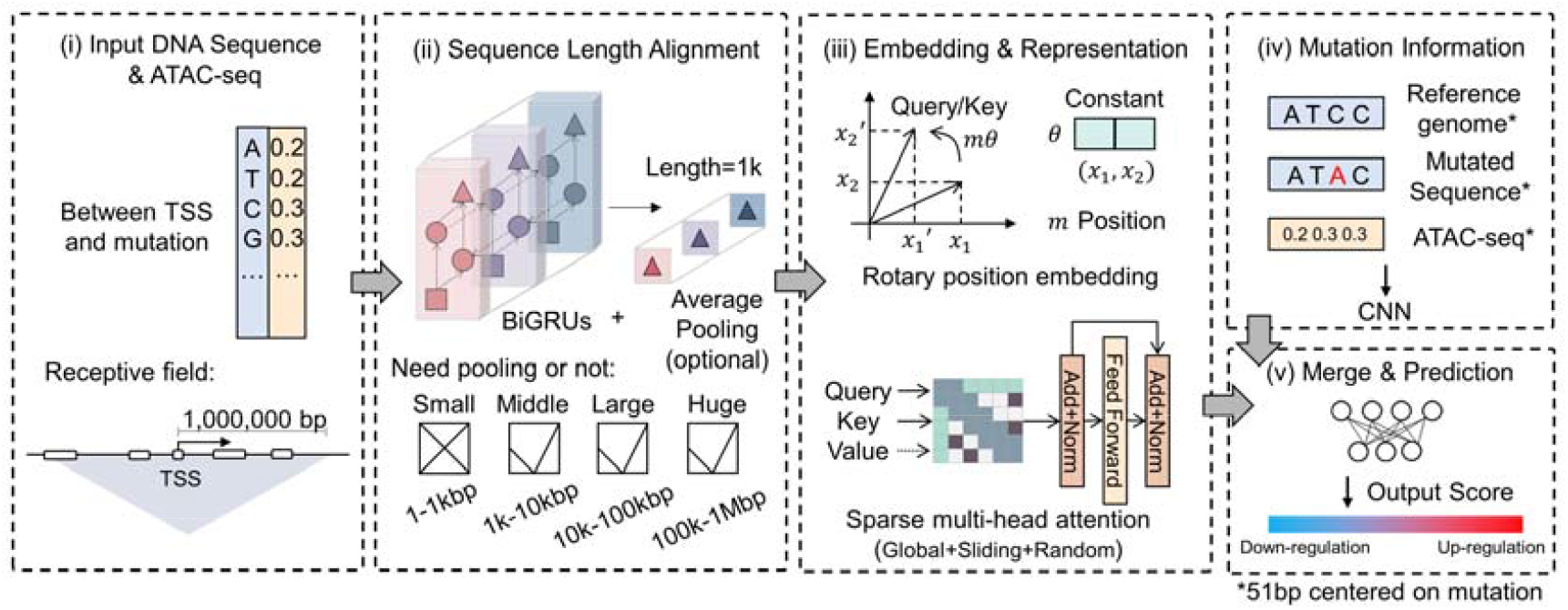
An overview of the EMO pipeline for non-coding mutation effect on eQTL sign prediction. The input of the model contains the ‘Between Branch’ and the ‘Mutation Branch’, One-Hot coding of the DNA sequence between the non-coding mutation and TSS, and the corresponding chromatin accessibility of the DNA sequence. The output is the predicted probabilities of a non-coding mutation affecting gene expression in an up-regulated manner.

To predict the trend of mutation effects on expression more precisely and economically on a larger scale, thus minimizing computational resources, we designed a ‘divide and conquer’ approach. This method categorizes mutations into four classes based on the distance from the variant to the TSS (the maximum length of DNA sequence input into the model), as shown in Figure 1.

For the input between the variant TSS and the variant itself, the encoded DNA sequence and the corresponding ATAC-seq are fed into a bidirectional Gated Recurrent Units (BiGRUs) module[18]. These BiGRUs can learn the sequence content from both past and future time steps[19]. Then, the average pooling[20] of varying sizes based on the variant TSS distance is applied. After this, we conduct rotational position encoding[21] to the pooled vectors and input them into sparse transformer layers that integrate different attention strategies[22] to further reduce the computational cost and keep the interpretability. For inputs centered on the variant, we use a CNN module. Finally, we merge all intermediate vectors and use the SoftMax function[23] for the final prediction output.

We trained EMO as four sub-models, each dedicated to handling non-coding mutations within different impact ranges (length of DNA sequence). These sub-models are referred to as ‘EMO-small’ (1bp-1kbp), ‘EMO-middle’ (1kbp-10kbp), ‘EMO-large’ (10kbp-100kbp), ‘EMO-huge’ (100kbp-1Mbp). The performance of these models was evaluated on the independent test sets, as illustrated in Figure 2(a)-(h) and Table 1. Despite a gradual decrease in the models’ capability to identify non-coding mutations affecting as ‘expression down-regulated’ (negative samples) with increasing sequence length (variant TSS distance), all sub-models exhibited an overall predictive AUC of 0.7 or higher. It’s worth noting that EMO’s predictions are not tissue-specific. As long as there is an ATAC-seq available for the DNA sequence, EMO can provide corresponding predictions, regardless of whether the tissue or cell to be predicted has appeared in the training set or not.

**Table 1.**
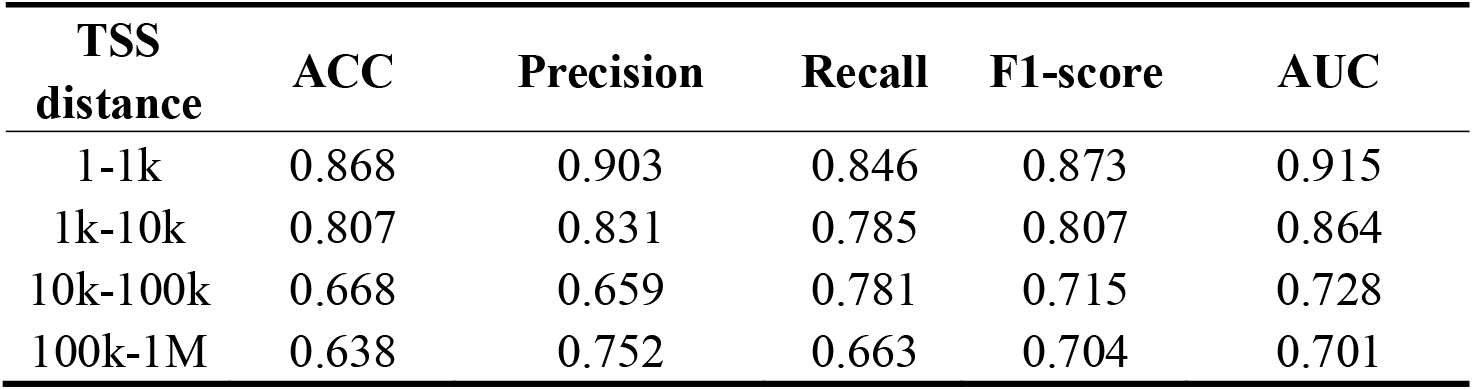
Model performance of EMO with different variant TSS distance.

| TSS distance | ACC | Precision | Recall | F1-score | AUC |
| --- | --- | --- | --- | --- | --- |
| 1-1k | 0.868 | 0.903 | 0.846 | 0.873 | 0.915 |
| 1k-10k | 0.807 | 0.831 | 0.785 | 0.807 | 0.864 |
| 10k-100k | 0.668 | 0.659 | 0.781 | 0.715 | 0.728 |
| 100k-1M | 0.638 | 0.752 | 0.663 | 0.704 | 0.701 |

**Figure 2.**
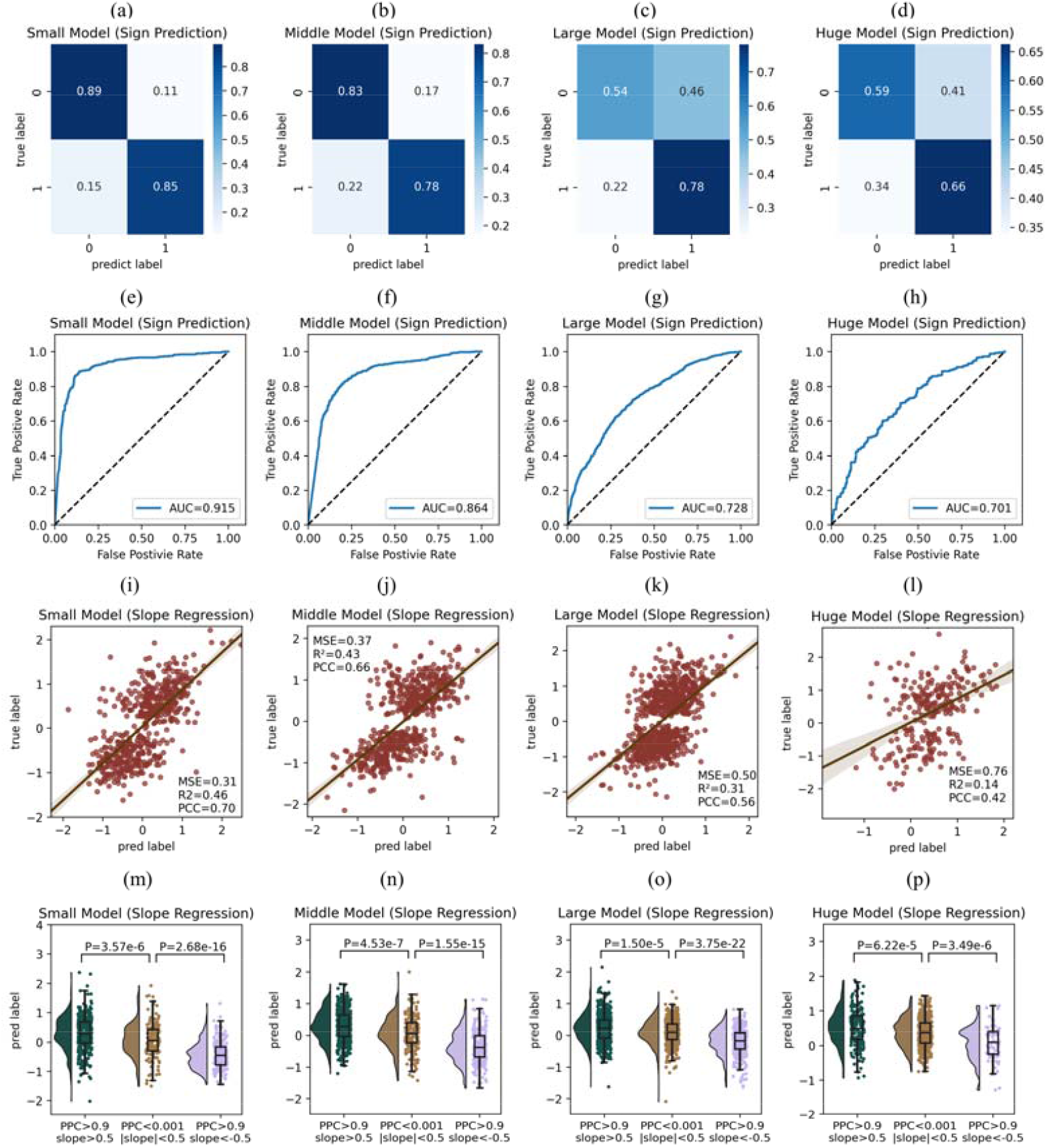
(a)-(h) Model performance of sub-models of EMO of varying sequence length (variant TSS distance) on the independent test set. The negative sample (label “0”) indicates that the mutation leads to a decrease in the expression of a downstream gene (down-regulation), whereas the positive sample (label “1”) signifies that the mutation results in an increased expression (up-regulation). (i)-(p) EMO analysis on employing slope regression to determine the magnitude of influence. The validation involved the creation of datasets comprising non-causal mutations and subsets representing up-regulation and down-regulation mutations, with significant differences observed in the distribution of predicted outcomes across these datasets using Mann-Whitney U test.

In addition to predicting the impact direction of non-coding mutations on downstream gene expression, EMO can also be trained to predict the specific magnitude of influence (using slope regression, depicted in Figure 2(i)-(l)). To validate the efficacy of the values derived from the EMO regression model, an additional dataset comprising non-causal mutations was constructed. This dataset consisted of non-coding mutations identified through fine-mapping with a posterior probability of being causal (PPC) less than 0.001 and an absolute slope value less than 0.5, indicative of mutations having no effect on downstream gene expression. Moreover, two subsets were extracted from the test dataset of EMO, encompassing mutations associated with up-regulation (PPC > 0.9, slope > 0.5) and down-regulation (PPC > 0.9, slope < -0.5) of gene expression, respectively. The EMO model underwent slope regression training on the training dataset and was subsequently evaluated on the aforementioned three datasets (Figure 2(m)-(p)). It is observed that there exists a significant distinction in the distribution of EMO predictions across these three datasets based on Mann-Whitney U test. Additionally, for the non-causal dataset, it was observed that the majority of predicted slope values fell within an absolute range of 1. This suggests that the slope values outputted by the EMO regression model to some extent reflect the degree of mutation impact, with mutations characterized by smaller absolute predicted slope values being recommended for classification as having negligible impact.

### 2.2 Representation visualization and ablation study

To investigate the representation ability of EMO and its capacity to extract high-level features from the input data, we used t-SNE[24] to visualize the initial features of the input branches and the final features prior to the output layer on the entire training dataset. As shown in Figure 3(a)-(h), no distinct or meaningful clusters were evident in the input features. However, after training, samples representing the two distinct categories exhibited a noticeable separation trend within the embedding space (Figure 3(i)-(l)), which became more pronounced in the case of short to medium DNA sequences, indicating that EMO could identify the proper classification hyperplane for the training dataset.

**Figure 3.**
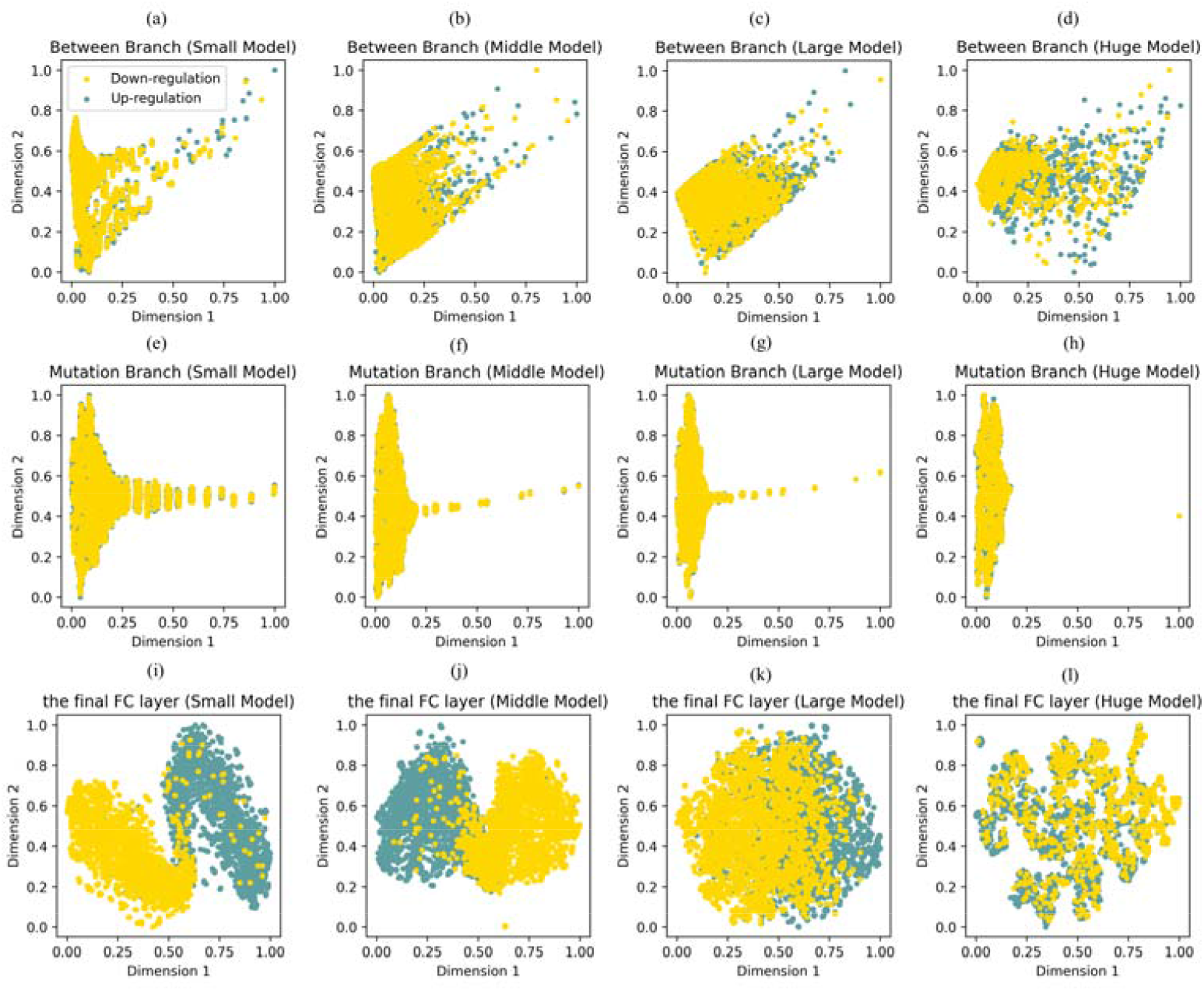
Representation visualization of EMO with input feature and the learned embedding before the output layer.

As the model comprised several processing modules, we conducted an ablation experiment to explore each module’s independent contributions to the model’s performance (Table 2). It was observed that the DNA sequence is undoubtedly the most crucial factor, while the ATAC-seq data, capable of reflecting tissue-specific information due to the tissue-specificity of eQTLs, is equally indispensable. Regarding the model architecture, we assessed the necessity of the architecture by replacing the original network layers with fully connected layers and found that the BiGRUs for capturing local information and the sparse self-attention layers for extracting long-range dependencies both made distinct individual contributions.

**Table 2.** Ablation study on the ‘EMO-Middle’ model.

| Component | Option | ACC | Precision | Recall | F1-score | AUC | Parameters |
| --- | --- | --- | --- | --- | --- | --- | --- |
| Input feature | Without ATAC-seq | 0.762 | 0.793 | 0.723 | 0.756 | 0.838 | 2,568,722 |
|  | Without DNA sequence | 0.589 | 0.623 | 0.494 | 0.551 | 0.619 | 2,494,994 |
| Local feature block | Without BiGRU* | 0.745 | 0.777 | 0.701 | 0.737 | 0.811 | 2,486,162 |
| Long-range block | Without attention layers* | 0.724 | 0.724 | 0.744 | 0.734 | 0.804 | 1,947,378 |
| EMO-Middle |  | **0.807** | **0.831** | **0.785** | **0.807** | **0.864** | 2,593,298 |

Numbers in bold indicate the best experimental results.
'\*' indicates that the component of EMO was replaced by fully connected layers, maintaining the same output dimensions.

### 2.3 Comparison between EMO and Enformer

We fine-tuned EMO for tissue specificity and compared the results with those from an end-to-end model trained independently on tissues (EMO-e2e) and Enformer[8] (Figure 4, Figure S4, and Table S5-8). This comparison was displayed across six example tissues with a large number of fine-mapped eQTLs to ensure that the different models could be sufficiently fitted. The eQTLs in the test set used for comparison were not included in EMO’s training set. For each eQTL, we obtained a 5313-dimensional output vector using the running interface officially released by Enformer. These output vectors were then used to train random forest models, varying by tissue. The implementation and the hyperparameters used for these random forest models were consistent with the reported descriptions[8].

**Figure 4.**
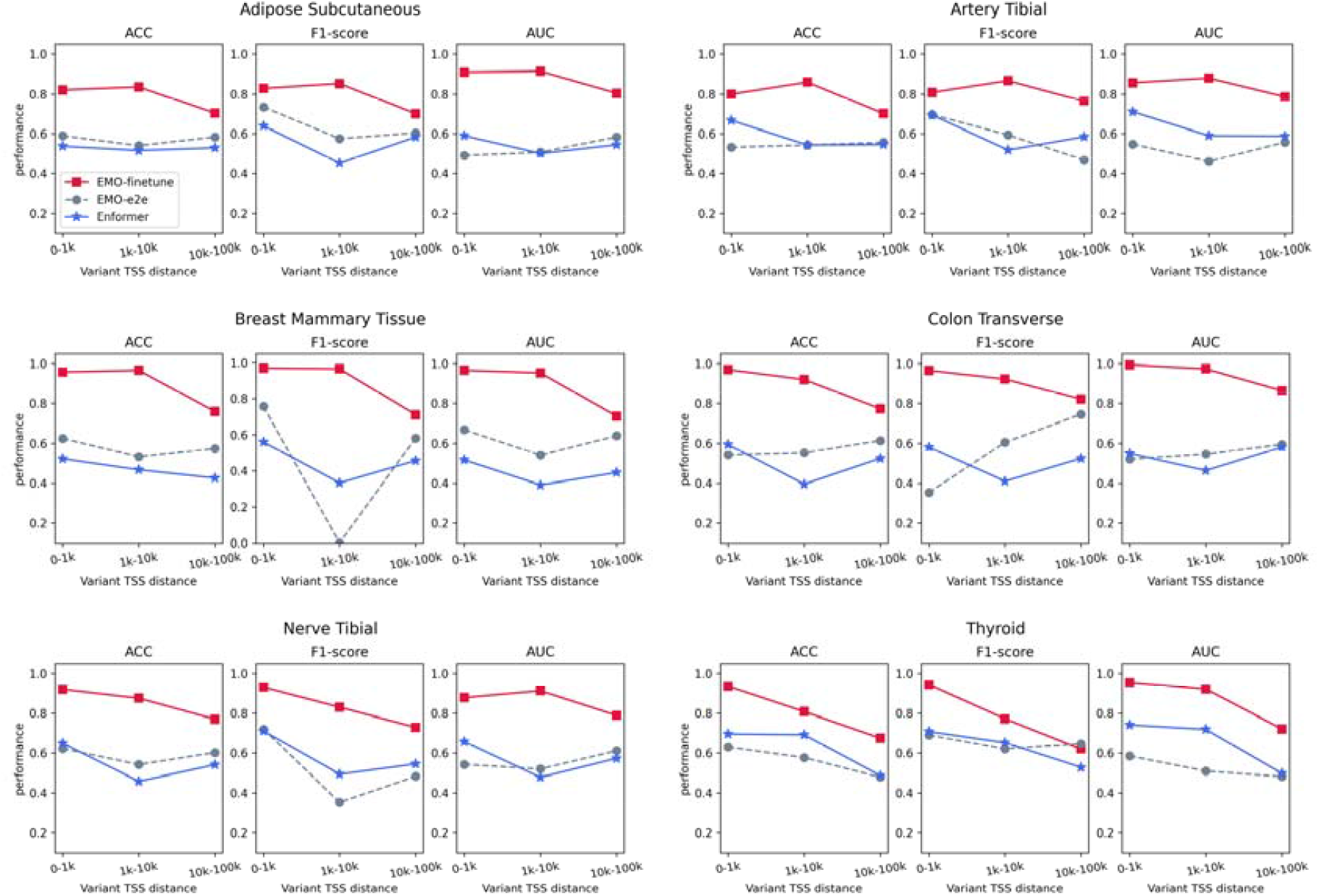
Performance comparison among EMO with tissue-specific fine-tuning, end-to-end training, and the retrained Enformer.

It is observed that after fine-tuning, EMO achieved better prediction performance on the tissues compared and showed stability (Figure 4), especially when the variant TSS is less than 1,000bp. The prediction performances of the ‘EMO-e2e’ model and Enformer were relatively similar. Notably, due to the relatively scarcity of eQTLs with TSS variant distances ranging between 1,001bp and 10,000bp of some tissues, the ‘EMO-e2e’ model and Enformer tend to experience a decrease in prediction accuracy in this condition, even worse than the performance of the model trained on longer TSS distances. This illustrates the impact of insufficient training data samples on end-to-end prediction. However, the ‘EMO-finetune’ model, leveraging embedded representations gained from pre-training on a larger sample volume, might carry information from the reference genome to the mutated DNA sequence that indicates the ‘direction of mutation’. This information might help more accurately determine the direction of the mutation’s effect on gene expression regulation, both up and down. Therefore, the ‘EMO-finetune’ model would achieve higher accuracy in tissue-specific predictions and, similarly, maintain consistent prediction performance across different prediction scenarios, as theorized by the difficulty of the predictions.

When comparing Enformer with the ‘EMO-e2e’ model, we found that Enformer exhibited greater stability. In some tissues, Enformer demonstrated superior prediction capability than the ‘EMO-e2e’ model (for example, in artery tibial and thyroid). This may suggest that functional information of variant (input of retrained Enformer) is more effective than the arrangement of bases (the DNA sequence itself, input of EMO) in end-to-end prediction. After fine-tuning, the EMO model showed improvements in both prediction performance and stability, suggesting that the pretraining might also allowed EMO to enhance potential representations capable of annotating the function of mutations.

### 2.4 Attention interpretability

Regulatory features like transcription factors and chromatin markers play vital roles in gene expression regulation and have been reported to be effective in associating eQTLs with target genes[25, 26]. In other words, in the sequence between non-coding mutations and the TSS, regulatory feature regions will provide more information for eQTLs sign prediction compared to other regions. Therefore, to investigate whether our model placed more emphasis on the information from such regulatory feature regions to enhance its accuracy, we map regulatory features to our input sequence and observe whether the attention weights of our model are significantly enriched in these regions.

We first obtained enhancer, TF binding, promotor, CTCF binding site, and open chromatin annotations from the Ensembl database[27] (version 2023-7-28) and mapped them to our training set input sequences. Then, we expanded the output attention weight vector to its original input length and separately aggregated the weights of regulatory feature regions and other regions. Due to the average length of regulatory feature regions being 950, sequences that are too short would be excessively annotated, and sequences that are too long would introduce larger errors due to the long pooling step. Here, we only conducted this attention interpretability part on middle and large-length sequences. The results of the Mann-Whitney one-tailed test[28] showed that regulatory feature regions had significantly greater attention weights than other regions in 75% of the heads (Table 3). This indicates that our model tends to focus more on the regulatory feature regions within the sequences, consistent with the previously mentioned relevance of regulatory features to the eQTL sign prediction task.

**Table 3.** Mann-Whitney one-tailed test p-values for attention weights in regulatory feature regions vs. other regions.

| Model | Head1 | Head2 | Head3 | Head4 |
| --- | --- | --- | --- | --- |
| Middle | 5.42e-09 | 2.40e-43 | 1 | 1.27e-37 |
| Large | 1 | 0 | 0 | 0 |

We also visualized two examples to further support the above conclusion (Figure 5). The eQTL for the “EMO-small” model is located at position 324,170 on chr11, from A to G, with the gene’s TSS downstream at position 5,530. The eQTL for the “EMO-large” model is located at position 15,194,525 on chr10, from A to G, with the gene’s TSS downstream at position 25,832. By comparing the attention input and attention output, we can observe that the model amplifies its focus on the regulatory feature region through the attention module.

**Figure 5.**
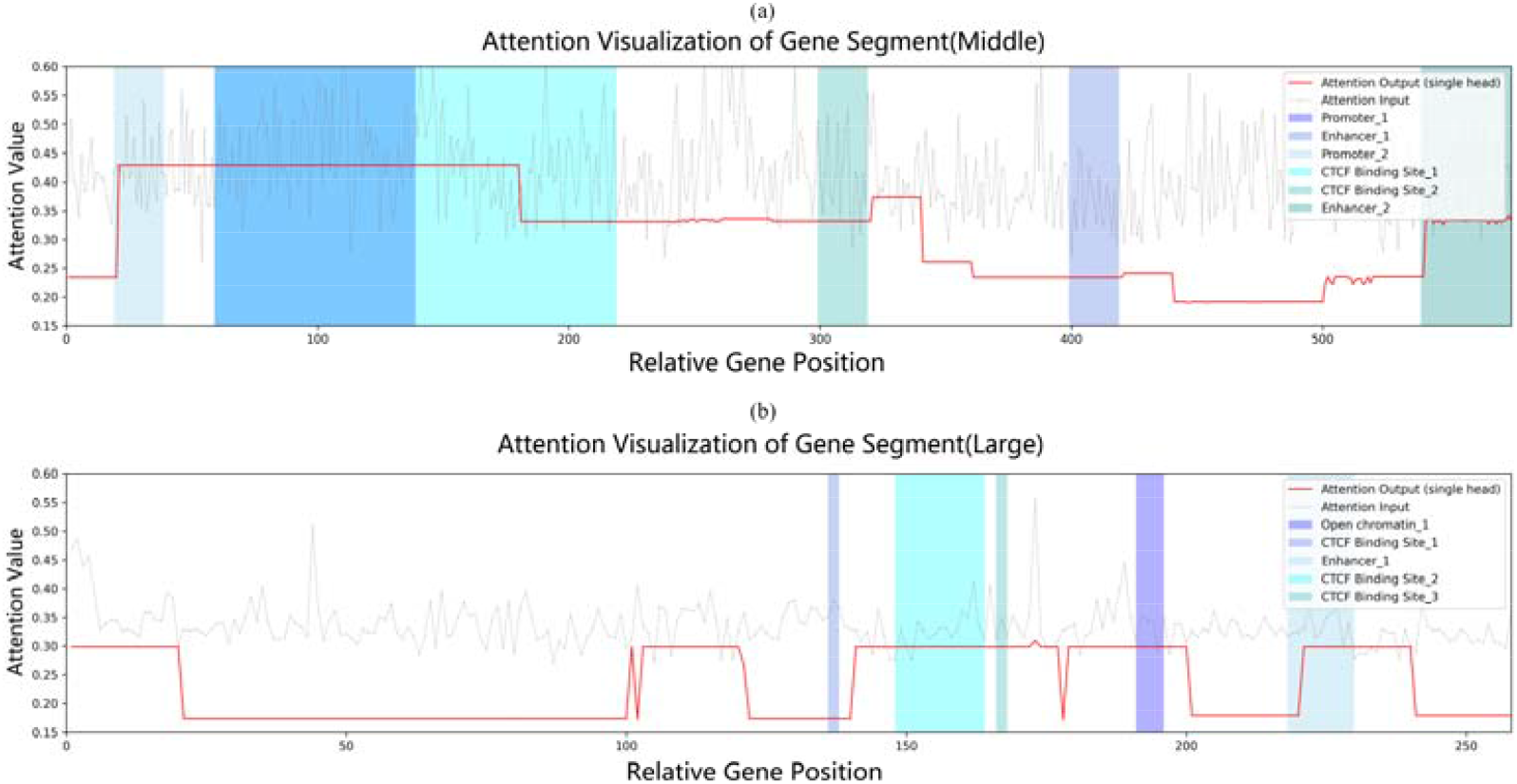
Visualization of attention IO and regulatory feature regions for two examples. Gray lines represent attention input; red lines represent attention output from head2. Colored segments denote annotated regulatory feature regions.

### 2.5 Tissue-specific fine-tuning on external brain eQTLs

As a tissue-agnostic universal pre-trained mode, EMO can seamlessly leverage tissue-specific ATAC-seq data and fine-tune it to various tissues and even migrate to single-cell resolution by learning the association between tissue-specific impact patterns of non-coding mutation and tissue ATAC-seq data.

To explore whether EMO can still perform stably when transferred to new tissues, we obtained the non-coding mutations from two brain tissues, the hippocampus and spinal cord, from MetaBrain[28], which were not included in the EMO training set. The hippocampus is critical for memory formation and spatial navigation, while the spinal cord is pivotal for transmitting information between the brain and the rest of the body[29, 30]. Understanding how genetic variations influence gene expression in these areas can provide insights into the complex processes governing brain development, neural connectivity, and synaptic plasticity.

As depicted in Figure 6, the fine-tuned EMO exhibited more reasonable predictions than the end-to-end predictions. For eQTLs with TSS distances ranging from 1,001bp to 10,000bp (trained using the ‘EMO-middle’ model), due to the small data sample size, models for the two tissues trained directly from end to end could not successfully fit and predicted all eQTLs in the test set as ‘Up-regulation’ (Figure 6(b), Figure 6(d)). However, models fine-tuned based on pre-trained embeddings achieved more reasonable prediction performance (Figure 6(a), Figure 6(c)). For eQTLs with TSS distances ranging from 10,001bp to 100,000bp (trained using the ‘EMO-large’ model), compared to the end-to-end models, fine-tuned EMO models showed improvements in the AUC on the eQTL test sets for the hippocampus and spinal cord by 0.164 and 0.079, respectively (Figure 6(e)-(h)). This finding suggests that the prior information contained in the embedding representation obtained from the pre-trained EMO contributes to better performance in predicting the up- and down-regulation of non-coding mutation effects for tissues with small sample sizes (Table S10-11).

**Figure 6.**
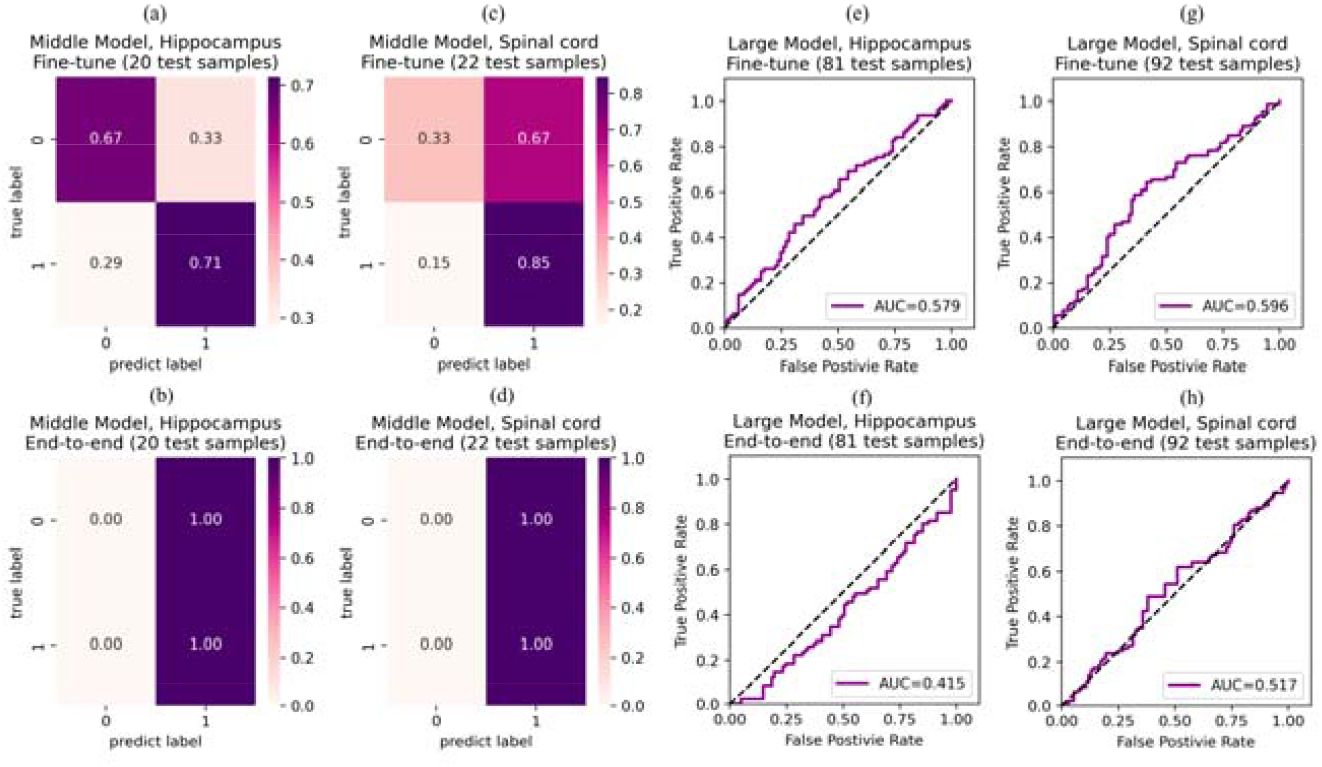
The prediction results of the fine-tuning and the end-to-end EMO model on eQTLs of two new tissues, hippocampus and spinal cord.

### 2.6 Transfer EMO to single-cell eQTLs

One of the core advantages of EMO is the ability to utilize transfer learning to apply pre-trained embeddings for predicting the up- and down-regulation effects of single-cell non-coding mutations. This advantage endows EMO with the capability to delve into personal transcriptome variations[14]. Also, many diseases, including cancer and autoimmune disorders, are driven by changes at the cellular level that can be obscured in bulk analyses[31, 32]. Single-cell eQTL studies can pinpoint the specific cell types and states where disease-associated genetic variants exert their effects, offering new avenues for targeted therapies and personalized medicine[33, 34].

Here, we collected the single-cell eQTL data of six immune cell types from the OneK1K cohort[35]. Each cell type underwent a separate fine-tuning process using EMO (as detailed in Table 4, Figure S6, and Table S12). It is observed that EMO demonstrated satisfactory performance across all cell types (AUC > 0.860), indicating the generalization potential of EMO’s pre-trained embedding representation in predicting the up- and down-regulation of non-coding mutation effects at the single-cell level. Such capability provides novel insights into the interpretation of cellular microenvironments, enhancing our understanding of cellular-scale genomic variations.

**Table 4.** The results of the transfer fine-tuned ‘EMO-large’ model on single-cell eQTLs of six cell types.

| Cell type | ACC | Precision | Recall | F1-score | AUC |
| --- | --- | --- | --- | --- | --- |
| B naïve | 0.815 | 0.882 | 0.600 | 0.714 | 0.861 |
| CD4 memory | 0.884 | 1.000 | 0.853 | 0.921 | 0.923 |
| CD4 naïve | 0.865 | 0.864 | 0.826 | 0.844 | 0.948 |
| CD8 memory | 0.756 | 0.714 | 0.870 | 0.784 | 0.862 |
| Dendritic cell | 0.818 | 0.893 | 0.893 | 0.893 | 0.919 |
| Natural killer cell | 0.823 | 0.913 | 0.677 | 0.778 | 0.911 |

### 2.7 Analysis of psychiatric disease-associated eQTL cases

Certain non-coding mutations, by altering the levels of gene expression, may increase or decrease an individual’s susceptibility to specific diseases[36]. To further investigate the performance of EMO on non-coding mutations associated with diseases, we collected two reported psychiatric disease-associated eQTLs as exploratory cases[28] (Table S13).

The A allele of variant rs4698412 was reported increased the expression of *CD38* and associated with increased risk for Parkinson’s Disease (PD), and rs1902660 was analyzed as an eQTL with higher expression of *TSPAN14* as putatively causal for Alzheimer’s Disease (AD). Both PD and AD are neurodegenerative disorders with complex etiologies[37]. By identifying eQTLs linked to these conditions, researchers can uncover the genetic variations that influence gene expression levels in the brain and other relevant tissues. This helps elucidate the molecular pathways and biological processes underlying the development and progression of these diseases.

We matched these two eQTLs with ATAC-seq data of the cerebellum[38, 39] and used EMO for prediction. As shown in Table S13, EMO correctly identified the direction of the effect of these two disease-associated non-coding mutations on gene expression, demonstrating its potential in deciphering the mechanisms of action at disease-risk SNPs.

## 3 Methods

### 3.1 Datasets

We curated cis-eQTL data of 48 tissue types in GTEx v8[40], retaining only those eQTL with a probability of being causal exceeding 0.9 following statistical fine-mapping[41]. Subsequently, we cross-referenced this eQTL dataset with 19 overlapping tissue-specific ATAC-seq data available in EpiMap[42] (Table S2-3), which were converted to hg38 human genome using CrossMap[43], and discarded the eQTLs with multi-site substitutions for subsequent prediction. The sequences between non-coding mutations and the TSS were extracted from the hg38 human genome. All eQTLs were classified into two categories, ‘up-regulation’ and ‘down-regulation’, based on their impact direction on the quantitative expression of the given gene. Due to the potentially biased model resulting from an overrepresentation of entries from the heart’s left ventricle and esophagus mucosa, we applied the down-sampling technique to these two tissues (Figure S1-3).

For each eQTL entry, we utilized the DNA sequence between non-coding mutation and TSS along with its corresponding ATAC-seq as the input features for deep learning. Specifically, in cases where multiple ATAC-seq results corresponded to the same tissue, we conducted a base-wise averaging:

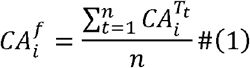

where 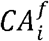represents the final chromatin accessibility of the *i-*th base in the gene sequence, *n* signifies the number of included ATAC-seq data for the current tissue within the dataset, and *CA*^*Tt*^ denotes the *t-*th of the *n* ATAC-seq data. Due to the broad range of the cis-eQTLs’ impact, extending from within 1,000 bp to as far as 1 million bp, blind data completeness for alignment purposes could potentially lead to unnecessary computational waste. Thus, based on the varying magnitudes of the extracted variant TSS distances, we partitioned the dataset into four distinct segments: the “small” dataset (1-1,000 bp), the “middle” dataset (1,001-10,000 bp), the “large” dataset (10,001-100,000 bp), and the “huge” dataset (100,001-1,000,000 bp). The training, validation, and test dataset were randomly distributed based on an 8:1:1 ratio (Table S1, Table S2, Figure S2).

We collected eQTLs of two new brain tissues, the hippocampus and spinal cord, from MetaBrain[28] for external validation (Table S9), and their corresponding tissue-specific ATAC-seq data was also downloaded from EpiMap (converted to hg38 with CrossMap), undergoing the same data preprocessing procedure. Additionally, to validate the transferability of EMO at the single-cell resolution, we acquired single-cell eQTLs from six cell types from the OneK1K cohort[35] and corresponding cell line ATAC-seq data from EpiMap (Table S3, Table S12).

### 3.2 Model architecture and prediction pipeline

We formulate the sign prediction of eQTLs as a 2-class classification problem, where each non-coding mutation that affected the gene expression may be assigned as either ‘up-regulation’ or ‘down-regulation’. EMO takes the One-Hot coding of the DNA sequence between the non-coding mutation and TSS, 51bp of the One-Hot coding of the DNA sequence of the reference genome centered on the mutation, 51bp of the One-Hot coding of the DNA sequence of the mutated sequence centered on the mutation, and the corresponding chromatin accessibility of the DNA sequence as input. The model’s output represents the predicted probabilities of a non-coding mutation affecting gene expression in an up-regulated manner.

EMO consists of five main stages (Figure 1): (i) encoding the DNA sequence between the non-coding mutation and TSS as ‘Between Branch,’ (ii) utilizing bidirectional Gated Recurrent Units (BiGRUs), aligning sequences of different lengths based on distinct-scale pooling strategies, (iii) applying rotary position embedding to the aligned features, subsequently inputting them into sparse multi-head attention layers, (iv) encoding the DNA sequence with a length of 51bp centered on the non-coding mutation site before and after the mutation, and inputting them into convolutional neural networks (CNN) as ‘Mutation Branch,’ (v) merging the feature representation from both the ‘Between Branch’ and the ‘Mutation Branch’ and utilizing fully-connected layers and the SoftMax function[23] for the final prediction, which are described in Sections 3.3, 3.4 and 3.5.

### 3.3 Alignment of DNA sequence with pooling

Since cis-eQTLs are SNPs within 1Mbp of a gene’s transcription start[13], aligning all DNA sequences to 1Mbp would result in computational waste for samples with medium to short DNA sequences. Therefore, we padded the input features with zeros based on the original length of DNA sequences: the sequences within the range of 1bp to 1kbp were padded to 1kbp, those within 1kbp to 10kbp were padded to 10kbp, and those within 10kbp to 100kbp were padded to 100kbp. Sequences longer than 100kbp were padded to 1Mbp.

After BiGRUs, we also conducted average pooling[20] for the sequences with lengths exceeding 1kbp:

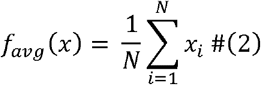

where *x* is a vector consisting of activation values from a rectangular area of *N* vectors in the dimension of the input features; keeping the embedding feature dimensions constant, we compressed the dimensions of the input sequences. The pooling size varied according to the magnitude of the length of the DNA sequences:

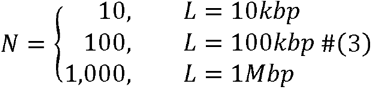

where *L* is the length of the input sequence after padding. Ultimately, the output feature of all DNA sequences was aligned to 1000 dimensions.

### 3.4 Representation learning with sparse Transformer

For the aligned 1000-dimensional output, we employed rotary position embedding[21] to indicate the positional information:

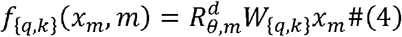

where *x*_*m*_ ∈ ℝ^*d*^ is the *d-*dimension feature embedding vector of the *m-*th token without position information, and 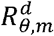 is the rotary matrix with pre-defined parameters *θ* ={*θ*_*I*_ = 10000^−2(*i*−1)/*d*^ *i* ∈ [1,2, …,*d/2*]}.

Then, we implemented sparse self-attention layers based on BigBird[22], which contains the global, sliding, and random multi-head self-attention for representation learning. The number of attention heads for each self-attention layer was set to 8.

### 3.5 Convolution and merge of mutation information

To preserve information from the reference genome near the mutation site and enhance the learning of mutation information with the network, we extracted reference genome and post-mutation DNA sequences with a total length of 51bp, centered around the mutation site, along with their corresponding ATAC-seq data. Then, this portion of the input as ‘Mutation Branch’ was fed into a convolutional module consisting of 2 CNN layers.

Then, we concatenated the embeddings from the ‘Between Branch’ and the ‘Mutation Branch’ and fed them into three fully connected network layers with decreasing numbers of artificial neurons. For the final output, we utilized the SoftMax function[23], resulting in the prediction outcome of EMO.

### 3.6 Details of comparison between EMO and Enformer

In comparing EMO with Enformer, we segmented EMO’s training set by tissue. We selected six example tissues with large numbers of fine-mapped eQTLs: adipose subcutaneous, artery tibial, breast mammary tissue, nerve tibial, testis, and thyroid, to ensure that each model had sufficient data for fitting. The test set used for comparison was also divided by tissue. Considering the different variant TSS distances, and given that Enformer lacks the capability to predict the effects of mutations with TSS distances greater than 100kb[8], we made comparisons at distances of 1-1kbp, 1k-10kbp, and 10k-100kbp.

For the ‘EMO-finetune’ model, we used the weights of the EMO model pre-trained on all training sets as the initial weights for the fine-tuning model, which was then retrained on tissue-specific training sets. For the ‘EMO-e2e’ model, we employed a random initialization method to conduct independent end-to-end training of EMO on the training sets divided by tissue. Regarding the Enformer model, for each eQTL, we obtained a 5,313-dimensional output vector from the officially released running interface, and trained a random forest model for each tissue. It has been reported that changing the hyperparameters of the random forest model negligibly affects model performance[8]. Therefore, we used the default hyperparameters of scikit-learn to implement these random forest models.

### 3.7 Evaluation metrics

As our study focuses on two-class classification for clinical drug response prediction, we evaluated the performance of our predictor using the following metrics[44]: accuracy, recall, precision, F1-score, and AUC, which were defined as:

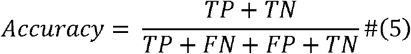

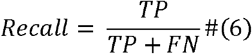

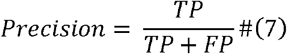

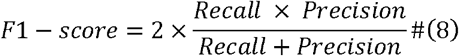

where TN, TP, FN, and FP represent the number of correctly predicted non-sensitive mutations, the number of correctly predicted sensitive mutations, the number of falsely predicted non-sensitive mutations, and the number of falsely predicted sensitive mutations, respectively. The AUC is computed using the trapezoidal rule to approximate the region under the receiver operating characteristic curve.

## 4 Discussion

An enduring challenge in regulatory genomics is the precise prediction of gene expression based solely on DNA sequence. Accurately predicting the sign of eQTLs can expedite the discovery of links between non-coding variants and phenotypic traits. In this research, we introduced EMO, a pre-trained model designed to predict both the up- and down-regulation effects of single non-coding mutations on gene expression. EMO can be fine-tuned to any tissue or cell type when ATAC-seq data is provided. We conducted a cost-effective exploration of long-range DNA sequence regulatory patterns, enhancing the interpretability of our findings. Our study offers a new insight into the impact of non-coding risk SNPs on the transcriptome, extending this understanding to single-cell resolution. Additionally, we validated EMO using multiple external datasets and demonstrated its potential in predicting disease-associated risk SNPs.

However, EMO has some limitations. In distinguishing between up-regulated and down-regulated non-coding mutations, EMO tends to make bolder predictions rather than conservative ones. This could lead to misjudgments and a failure to recognize noncausal variants. Though we have presented an alternative approach, slope regression, to preemptively determine which mutations may potentially be causal, a more prudent approach involves employing EMO on established risk SNPs to facilitate a deeper understanding of the mechanistic impact of SNPs on gene expression. Moreover, EMO’s predictive performance for medium to long DNA sequences could be further improved.

Looking ahead, we aim to continue refining EMO, enabling it to rapidly annotate genes affected by specific risk non-coding mutations on a large scale and indicate the direction of such effects. As large language pre-trained models advance, we also plan to explore incorporating semantic embeddings of DNA sequences to optimize EMO’s performance.

## Code availability

The data and source code of EMO is available at https://github.com/Liuzhe30/EMO, with the running interface of sign prediction and slope regression provided.

## References

1. Zhou, J., et al., Deep learning sequence-based ab initio prediction of variant effects on expression and disease risk. Nature genetics, 2018. 50(8): p. 1171–1179.

2. Ashley, E.A., Towards precision medicine. Nature Reviews Genetics, 2016. 17(9): p. 507–522.

3. Aguet, F., et al., Genetic effects on gene expression across human tissues. Nature, 2017. 550(7675): p. 204–213.

4. Cooper, G.M. and J. Shendure, Needles in stacks of needles: finding disease-causal variants in a wealth of genomic data. Nature Reviews Genetics, 2011. 12(9): p. 628–640.

5. Lek, M., et al., Analysis of protein-coding genetic variation in 60,706 humans. Nature, 2016. 536(7616): p. 285–291.

6. Consortium, G.P., A map of human genome variation from population scale sequencing. Nature, 2010. 467(7319): p. 1061.

7. Khurana, E., et al., Role of non-coding sequence variants in cancer. Nature Reviews Genetics, 2016. 17(2): p. 93–108.

8. Avsec, Ž., et al., Effective gene expression prediction from sequence by integrating long-range interactions. Nature methods, 2021. 18(10): p. 1196–1203.

9. French, J.D. and S.L. Edwards, The Role of Noncoding Variants in Heritable Disease. Trends in Genetics, 2020. 36(11): p. 880–891.

10. Zeng, H. and D.K. Gifford, Predicting the impact of non-coding variants on DNA methylation. Nucleic Acids Research, 2017. 45(11): p. e99–e99.

11. Zhou, J. and O.G. Troyanskaya, Predicting effects of noncoding variants with deep learning–based sequence model. Nature methods, 2015. 12(10): p. 931–934.

12. Kelley, D.R., Cross-species regulatory sequence activity prediction. PLoS computational biology, 2020. 16(7): p. e1008050.

13. Gaynor, S.M., et al., Connectivity in eQTL networks dictates reproducibility and genomic properties. Cell Rep Methods, 2022. 2(5): p. 100218.

14. Huang, C., et al., Personal transcriptome variation is poorly explained by current genomic deep learning models. Nature Genetics, 2023. 55(12): p. 2056–2059.

15. Sasse, A., et al., Benchmarking of deep neural networks for predicting personal gene expression from DNA sequence highlights shortcomings. Nature Genetics, 2023. 55(12): p. 2060–2064.

16. Musunuru, K., et al., From noncoding variant to phenotype via SORT1 at the 1p13 cholesterol locus. Nature, 2010. 466(7307): p. 714–719.

17. Painter, J.N., et al., A Common Variant at the 14q32 Endometrial Cancer Risk Locus Activates AKT1 through YY1 Binding. Am J Hum Genet, 2016. 98(6): p. 1159–1169.

18. Tham, M., BIDIRECTIONAL GATED RECURRENT UNIT FOR SHALLOW PARSING. Indian Journal of Computer Science and Engineering, 2020. 11: p. 517–521.

19. Tran, T.U., H.T.T. Hoang, and H.X. Huynh. Aspect Extraction with Bidirectional GRU and CRF. in 2019 IEEE-RIVF International Conference on Computing and Communication Technologies (RIVF). 2019.

20. Sharma, S. and D.R. Mehra, Implications of Pooling Strategies in Convolutional Neural Networks: A Deep Insight. Foundations of Computing and Decision Sciences, 2019. 44: p. 303–330.

21. Su, J., et al., Roformer: Enhanced transformer with rotary position embedding. arXiv preprint arXiv:2104.09864, 2021.

22. Zaheer, M., et al., Big bird: Transformers for longer sequences. Advances in neural information processing systems, 2020. 33: p. 17283–17297.

23. Gibbs, J.W., Elementary principles in statistical mechanics: developed with especial reference to the rational foundations of thermodynamics. 1902: C. Scribner’s sons.

24. van der Maaten, L. and G. Hinton, Viualizing data using t-SNE. Journal of Machine Learning Research, 2008. 9: p. 2579–2605.

25. Wang, D., A. Rendon, and L. Wernisch, Transcription factor and chromatin features predict genes associated with eQTLs. Nucleic Acids Research, 2012. 41(3): p. 1450–1463.

26. Flynn, E.D., et al., Transcription factor regulation of eQTL activity across individuals and tissues. Cold Spring Harbor Laboratory, 2021.

27. Cunningham, F., et al., Ensembl 2022. Nucleic Acids Research, 2021. 50(D1): p. D988–D995.

28. de Klein, N., et al., Brain expression quantitative trait locus and network analyses reveal downstream effects and putative drivers for brain-related diseases. Nature Genetics, 2023. 55(3): p. 377–388.

29. Eichenbaum, H., The role of the hippocampus in navigation is memory. J Neurophysiol, 2017. 117(4): p. 1785–1796.

30. Kakulas, B.A., Neuropathology: the foundation for new treatments in spinal cord injury. Spinal Cord, 2004. 42(10): p. 549–563.

31. Bolon, B., Cellular and molecular mechanisms of autoimmune disease. Toxicol Pathol, 2012. 40(2): p. 216–29.

32. Rahat, M.A. and J. Shakya, Parallel Aspects of the Microenvironment in Cancer and Autoimmune Disease. Mediators Inflamm, 2016. 2016: p. 4375120.

33. Fujita, M., et al., Cell subtype-specific effects of genetic variation in the Alzheimer’s disease brain. Nature Genetics, 2024.

34. Levinsohn, J., et al., Combing Genome-Wide Association Studies and Single-Cell Analysis to Elucidate the Mechanisms of Kidney Disease: Proceedings of the Henry Shavelle Professorship. Glomerular Dis, 2023. 3(1): p. 258–265.

35. Yazar, S., et al., Single-cell eQTL mapping identifies cell type–specific genetic control of autoimmune disease. Science, 2022. 376(6589): p. eabf3041.

36. Knight, J.C., Functional implications of genetic variation in non-coding DNA for disease susceptibility and gene regulation. Clin Sci (Lond), 2003. 104(5): p. 493–501.

37. Chin-Chan, M., J. Navarro-Yepes, and B. Quintanilla-Vega, Environmental pollutants as risk factors for neurodegenerative disorders: Alzheimer and Parkinson diseases. Front Cell Neurosci, 2015. 9: p. 124.

38. Wu, T. and M. Hallett, The cerebellum in Parkinson’s disease. Brain, 2013. 136(Pt 3): p. 696–709.

39. Jacobs, H.I.L., et al., The cerebellum in Alzheimer’s disease: evaluating its role in cognitive decline. Brain, 2018. 141(1): p. 37–47.

40. Consortium, G., The GTEx Consortium atlas of genetic regulatory effects across human tissues. Science, 2020. 369(6509): p. 1318–1330.

41. Wang, Q.S., et al., Leveraging supervised learning for functionally informed fine-mapping of cis-eQTLs identifies an additional 20,913 putative causal eQTLs. Nature Communications, 2021. 12(1): p. 3394.

42. Boix, C.A., et al., Regulatory genomic circuitry of human disease loci by integrative epigenomics. Nature, 2021. 590(7845): p. 300–307.

43. Zhao, H., et al., CrossMap: a versatile tool for coordinate conversion between genome assemblies. Bioinformatics, 2014. 30(7): p. 1006–7.

44. Salavati, H., et al., Drug transport modeling in solid tumors: A computational exploration of spatial heterogeneity of biophysical properties. Computers in Biology and Medicine, 2023. 163.

