## Supplementary Materials for "EMO: Predicting Non-coding Mutation-induced Up- and Down-regulation of Risk Gene Expression using Deep Learning"

**Tables**

Table S1. The final distribution of different variant TSS distance of the EMO dataset after post data processing, with the samples count after the dataset partitioning. In datasets with different TSS distance, the distribution of positive and negative samples is uniform in both the training set and the test set. During model training, one-ninth of the training data will be re-divided into validation data.

| **Model** | **variant TSS distance** | **Train** | **Test** | **Up-regulation** | | | **Down-regulation** | | | **All** |
| --- | --- | --- | --- | --- | --- | --- | --- | --- | --- | --- |
|  |  |  |  | **Train** | **Test** | **All** | **Train** | **Test** | **All** |  |
| Small | 1-  1,000 | 4,704 | 551 | 2,462 | 298 | 2,760 | 2,242 | 253 | 2,495 | 5,255 |
| Middle | 1,001-10,000 | 5,752 | 696 | 2,922 | 358 | 3,280 | 2,830 | 338 | 3,168 | 6,448 |
| Large | 10,001-100,000 | 9,029 | 943 | 4,684 | 502 | 5,186 | 4,345 | 441 | 4,786 | 9,972 |
| Huge | 100,001-1,000,000 | 2,461 | 246 | 1,455 | 160 | 1,615 | 1,006 | 86 | 1,092 | 2,707 |
| All | | 21,946 | 2,436 | 12,841 | | | 11,541 | | | 24,382 |

Table S2. The distribution of different tissues of the EMO dataset after partitioning. During model training, one-ninth of the training data will be re-divided into validation data.

| **Tissue** | **Train** | **Test** | **Tissue** | **Train** | **Test** |
| --- | --- | --- | --- | --- | --- |
| Adipose subcutaneous | 2,034 | 226 | Liver | 455 | 51 |
| Adrenal gland | 681 | 76 | Nerve tibial | 2,324 | 259 |
| Artery tibial | 2,031 | 226 | Ovary | 467 | 52 |
| Brain cerebellum | 1,109 | 124 | Pancreas | 1,020 | 114 |
| Brain cortex | 740 | 83 | Prostate | 609 | 68 |
| Brest mammary tissue | 1,115 | 124 | Spleen | 984 | 110 |
| Colon sigmoid | 990 | 111 | Stomach | 829 | 93 |
| Colon transverse | 1,137 | 127 | Testis | 1,791 | 200 |
| Esophagus mucosa | 1,221 | 136 | Thyroid | 2,286 | 254 |
| Heart left ventricle | 1,216 | 136 |  |  |  |

Table S3. The fetch number of ATAC-seq data downloaded from EpiMap.

| **Tissue** | **BSS ID** | **Tissue** | **BSS ID** |
| --- | --- | --- | --- |
| Adipose subcutaneous | BSS01671 | Esophagus mucosa | BSS00323 |
| Adrenal gland | BSS00047 |  | BSS00324 |
|  | BSS00055 | Heart left ventricle | BSS00506 |
|  | BSS00056 |  | BSS00507 |
| Artery tibial | BSS01839 | Liver | BSS01519 |
| Brain cerebellum | BSS00205 | Nerve tibial | BSS01840 |
| Brain cortex | BSS01452 | Ovary | BSS01402 |
| Brest mammary tissue | BSS00145 | Pancreas | BSS00121 |
|  | BSS00146 |  | BSS00122 |
|  | BSS00148 |  | BSS00123 |
| Colon sigmoid | BSS01542 |  | BSS00124 |
|  | BSS01543 | Prostate | BSS01457 |
|  | BSS01546 | Spleen | BSS01629 |
|  | BSS01547 | Stomach | BSS01639 |
| Colon transverse | BSS01848 |  | BSS01653 |
|  | BSS01849 | Testis | BSS01715 |
|  | BSS01850 | Thyroid | BSS01831 |
|  | BSS01851 |  | BSS01832 |
|  |  |  | BSS01835 |

Table S4. Training and model architecture details.

| **Model** | **Variant TSS distance** | **No. of BiGRU units** | **No. of Transformer layers** | **Attention head(s) per Transformer layer** | **GPU** | **Batch size** | **No. of model hypermeters** |
| --- | --- | --- | --- | --- | --- | --- | --- |
| Small | 1-1k | 32 | 2 | 4 | 1*NVIDIA 3090 | 32 | 1,419,602 |
| Middle | 1k-10k | 64 | 2 | 4 | 1*NVIDIA 3090 | 32 | 2,593,298 |
| Large | 10k-100k | 64 | 2 | 4 | 1*NVIDIA 3090 | 16 | 8,353,298 |
| Huge | 100k-1M | 32 | 2 | 4 | 2*NVIDIA L20 | 2 | 65,355,602 |

Table S5. Tissue-specific fine-tune performance of “EMO-small” model.

| **Option** | **Tissue** | **ACC** | **Precision** | **Recall** | **F1-score** | **AUC** |
| --- | --- | --- | --- | --- | --- | --- |
| Fine-tuning | Adipose subcutaneous | 0.821 | 0.895 | 0.773 | 0.829 | 0.909 |
|  | Artery tibial | 0.800 | 0.826 | 0.792 | 0.809 | 0.857 |
|  | Brest mammary tissue | 0.958 | 0.941 | 1.000 | 0.970 | 0.967 |
|  | Colon transverse | 0.970 | 0.933 | 1.000 | 0.966 | 0.994 |
|  | Nerve tibial | 0.919 | 0.909 | 0.952 | 0.930 | 0.878 |
|  | Testis | 0.761 | 0.800 | 0.769 | 0.784 | 0.792 |
|  | Thyroid | 0.935 | 0.926 | 0.962 | 0.943 | 0.954 |
| End-to-end | Adipose subcutaneous | 0.590 | 0.579 | 1.000 | 0.733 | 0.492 |
|  | Artery tibial | 0.533 | 0.533 | 1.000 | 0.696 | 0.548 |
|  | Brest mammary tissue | 0.625 | 0.667 | 0.875 | 0.757 | 0.668 |
|  | Colon transverse | 0.542 | 0.360 | 0.346 | 0.353 | 0.520 |
|  | Nerve tibial | 0.622 | 0.621 | 0.857 | 0.720 | 0.543 |
|  | Testis | 0.608 | 0.643 | 0.692 | 0.667 | 0.556 |
|  | Thyroid | 0.630 | 0.655 | 0.731 | 0.691 | 0.585 |
| EMO prediction | Adipose subcutaneous | 0.821 | 0.857 | 0.818 | 0.837 | 0.902 |
|  | Artery tibial | 0.800 | 0.826 | 0.792 | 0.809 | 0.849 |
|  | Brest mammary tissue | 0.958 | 0.951 | 1.000 | 0.970 | 0.966 |
|  | Colon transverse | 0.970 | 0.933 | 1.000 | 0.966 | 0.981 |
|  | Nerve tibial | 0.892 | 0.870 | 0.952 | 0.909 | 0.888 |
|  | Testis | 0.717 | 0.710 | 0.846 | 0.772 | 0.788 |
|  | Thyroid | 0.935 | 0.926 | 0.962 | 0.943 | 0.956 |

Table S6. Tissue-specific fine-tune performance of “EMO-middle” model.

| **Option** | **Tissue** | **ACC** | **Precision** | **Recall** | **F1-score** | **AUC** |
| --- | --- | --- | --- | --- | --- | --- |
| Fine-tuning | Adipose subcutaneous | 0.836 | 0.853 | 0.853 | 0.853 | 0.914 |
|  | Artery tibial | 0.860 | 0.839 | 0.897 | 0.867 | 0.880 |
|  | Brain cerebellum | 0.743 | 0.722 | 0.765 | 0.743 | 0.839 |
|  | Brest mammary tissue | 0.967 | 0.933 | 1.000 | 0.966 | 0.954 |
|  | Colon sigmoid | 0.967 | 0.933 | 1.000 | 0.966 | 0.989 |
|  | Colon transverse | 0.921 | 1.000 | 0.857 | 0.923 | 0.974 |
|  | Nerve tibial | 0.875 | 0.815 | 0.846 | 0.830 | 0.912 |
|  | Pancreas | 0.829 | 0.800 | 0.800 | 0.800 | 0.907 |
|  | Spleen | 0.821 | 0.882 | 0.833 | 0.857 | 0.828 |
|  | Thyroid | 0.808 | 0.772 | 0.772 | 0.772 | 0.921 |
| End-to-end | Adipose subcutaneous | 0.541 | 0.594 | 0.559 | 0.576 | 0.508 |
|  | Artery tibial | 0.544 | 0.543 | 0.655 | 0.594 | 0.462 |
|  | Brain cerebellum | 0.514 | 0.000 | 0.000 | 0.000 | 0.504 |
|  | Brest mammary tissue | 0.533 | 0.000 | 0.000 | 0.000 | 0.541 |
|  | Colon sigmoid | 0.720 | 0.800 | 0.750 | 0.774 | 0.758 |
|  | Colon transverse | 0.553 | 0.591 | 0.619 | 0.605 | 0.546 |
|  | Nerve tibial | 0.542 | 0.360 | 0.346 | 0.353 | 0.520 |
|  | Pancreas | 0.571 | 0.500 | 0.400 | 0.444 | 0.596 |
|  | Spleen | 0.607 | 0.667 | 0.778 | 0.718 | 0.592 |
|  | Thyroid | 0.577 | 0.500 | 0.818 | 0.621 | 0.510 |
| EMO prediction | Adipose subcutaneous | 0.853 | 0.857 | 0.882 | 0.870 | 0.908 |
|  | Artery tibial | 0.842 | 0.833 | 0.862 | 0.847 | 0.873 |
|  | Brain cerebellum | 0.771 | 0.737 | 0.824 | 0.777 | 0.825 |
|  | Brest mammary tissue | 0.967 | 0.933 | 1.000 | 0.966 | 0.946 |
|  | Colon sigmoid | 0.960 | 1.000 | 0.938 | 0.968 | 0.976 |
|  | Colon transverse | 0.921 | 1.000 | 0.857 | 0.923 | 0.970 |
|  | Nerve tibial | 0.875 | 0.774 | 0.923 | 0.842 | 0.897 |
|  | Pancreas | 0.800 | 0.786 | 0.733 | 0.759 | 0.906 |
|  | Spleen | 0.786 | 0.833 | 0.833 | 0.833 | 0.808 |
|  | Thyroid | 0.827 | 0.810 | 0.773 | 0.791 | 0.923 |

Table S7. Tissue-specific fine-tune performance of “EMO-large” model.

| **Option** | **Tissue** | **ACC** | **Precision** | **Recall** | **F1-score** | **AUC** |
| --- | --- | --- | --- | --- | --- | --- |
| Fine-tuning | Adipose subcutaneous | 0.703 | 0.660 | 0.745 | 0.700 | 0.805 |
|  | Artery tibial | 0.701 | 0.671 | 0.887 | 0.764 | 0.787 |
|  | Brain cerebellum | 0.667 | 0.684 | 0.813 | 0.743 | 0.741 |
|  | Brain cortex | 0.688 | 0.765 | 0.684 | 0.722 | 0.684 |
|  | Brest mammary tissue | 0.759 | 0.696 | 0.727 | 0.711 | 0.736 |
|  | Colon sigmoid | 0.773 | 0.783 | 0.783 | 0.783 | 0.846 |
|  | Colon transverse | 0.773 | 0.742 | 0.920 | 0.821 | 0.865 |
|  | Nerve tibial | 0.770 | 0.833 | 0.648 | 0.729 | 0.789 |
|  | Pancreas | 0.587 | 0.500 | 0.631 | 0.558 | 0.678 |
|  | Spleen | 0.771 | 0.774 | 0.857 | 0.814 | 0.855 |
|  | Stomach | 0.706 | 0.696 | 0.842 | 0.762 | 0.798 |
|  | Testis | 0.592 | 0.610 | 0.625 | 0.617 | 0.677 |
|  | Thyroid | 0.675 | 0.705 | 0.554 | 0.620 | 0.721 |
| End-to-end | Adipose subcutaneous | 0.584 | 0.542 | 0.681 | 0.604 | 0.584 |
|  | Artery tibial | 0.557 | 0.679 | 0.358 | 0.469 | 0.558 |
|  | Brain cerebellum | 0.574 | 0.636 | 0.656 | 0.646 | 0.593 |
|  | Brain cortex | 0.406 | 0.000 | 0.000 | 0.000 | 0.415 |
|  | Brest mammary tissue | 0.574 | 0.485 | 0.727 | 0.582 | 0.638 |
|  | Colon sigmoid | 0.523 | 0.523 | 1.000 | 0.687 | 0.590 |
|  | Colon transverse | 0.614 | 0.595 | 1.000 | 0.746 | 0.595 |
|  | Nerve tibial | 0.602 | 0.627 | 0.389 | 0.483 | 0.611 |
|  | Pancreas | 0.587 | 0.000 | 0.000 | 0.000 | 0.607 |
|  | Spleen | 0.521 | 0.727 | 0.286 | 0.410 | 0.463 |
|  | Stomach | 0.588 | 0.600 | 0.789 | 0.682 | 0.626 |
|  | Testis | 0.434 | 0.429 | 0.225 | 0.295 | 0.462 |
|  | Thyroid | 0.479 | 0.479 | 1.000 | 0.647 | 0.482 |
| EMO prediction | Adipose subcutaneous | 0.653 | 0.588 | 0.851 | 0.696 | 0.738 |
|  | Artery tibial | 0.701 | 0.688 | 0.830 | 0.752 | 0.786 |
|  | Brain cerebellum | 0.667 | 0.684 | 0.813 | 0.743 | 0.712 |
|  | Brain cortex | 0.594 | 0.636 | 0.737 | 0.683 | 0.641 |
|  | Brest mammary tissue | 0.685 | 0.586 | 0.773 | 0.667 | 0.726 |
|  | Colon sigmoid | 0.727 | 0.739 | 0.739 | 0.739 | 0.801 |
|  | Colon transverse | 0.818 | 0.774 | 0.960 | 0.857 | 0.840 |
|  | Nerve tibial | 0.690 | 0.630 | 0.852 | 0.724 | 0.749 |
|  | Pancreas | 0.500 | 0.429 | 0.632 | 0.511 | 0.635 |
|  | Spleen | 0.792 | 0.800 | 0.857 | 0.828 | 0.847 |
|  | Stomach | 0.676 | 0.682 | 0.789 | 0.732 | 0.753 |
|  | Testis | 0.579 | 0.587 | 0.675 | 0.628 | 0.659 |
|  | Thyroid | 0.658 | 0.621 | 0.732 | 0.672 | 0.714 |

Table S8. Tissue-specific performance of Enformer-trained random forest with different sequence length, with DNA sequence before and after mutation as input.

| **Tissue** | **Variant TSS distance** | **ACC** | **Precision** | **Recall** | **F1-score** | **AUC** |
| --- | --- | --- | --- | --- | --- | --- |
| Adipose subcutaneous | 1-1k | 0.538 | 0.571 | 0.727 | 0.640 | 0.590 |
|  | 1k-10k | 0.517 | 0.600 | 0.364 | 0.453 | 0.502 |
|  | 10k-100k | 0.530 | 0.500 | 0.702 | 0.584 | 0.545 |
| Artery tibial | 1-1k | 0.667 | 0.680 | 0.708 | 0.694 | 0.710 |
|  | 1k-10k | 0.544 | 0.560 | 0.483 | 0.519 | 0.590 |
|  | 10k-100k | 0.546 | 0.585 | 0.585 | 0.585 | 0.588 |
| Colon transverse | 1-1k | 0.594 | 0.500 | 0.692 | 0.581 | 0.549 |
|  | 1k-10k | 0.395 | 0.444 | 0.381 | 0.410 | 0.463 |
|  | 10k-100k | 0.524 | 0.647 | 0.440 | 0.524 | 0.581 |
| Nerve tibial | 1-1k | 0.649 | 0.667 | 0.762 | 0.711 | 0.657 |
|  | 1k-10k | 0.458 | 0.373 | 0.731 | 0.494 | 0.478 |
|  | 10k-100k | 0.541 | 0.508 | 0.588 | 0.545 | 0.573 |
| Testis | 1-1k | 0.630 | 0.696 | 0.615 | 0.653 | 0.688 |
|  | 1k-10k | 0.537 | 0.621 | 0.562 | 0.590 | 0.544 |
|  | 10k-100k | 0.579 | 0.580 | 0.725 | 0.644 | 0.634 |
| Thyroid | 1-1k | 0.696 | 0.773 | 0.654 | 0.708 | 0.741 |
|  | 1k-10k | 0.692 | 0.625 | 0.682 | 0.652 | 0.719 |
|  | 10k-100k | 0.487 | 0.465 | 0.611 | 0.528 | 0.498 |

Table S9. New brain eQTLs collected from the MetaBrain database.

| **Tissue** | **Small eQTLs** | **Middle eQTLs** | **Large eQTLs** | **Huge eQTLs** |
| --- | --- | --- | --- | --- |
| Hippocampus | 44 | 193 | 804 | 3,151 |
| Spinal cord | 60 | 217 | 913 | 3,754 |

Table S10. New brain eQTLs evaluation from the MetaBrain database on “EMO-middle” model.

| **Tissue** | **Option** | **ACC** | **Precision** | **Recall** | **F1-score** | **AUC** |
| --- | --- | --- | --- | --- | --- | --- |
| Hippocampus | Fine-tune | 0.700 | 0.833 | 0.714 | 0.769 | 0.765 |
|  | End-to-end | - | - | - | - | - |
| Spinal cord | Fine-tune | 0.636 | 0.647 | 0.846 | 0.733 | 0.579 |
|  | End-to-end | - | - | - | - | - |

“-” indicates that the model cannot fit data.

Table S11. New brain eQTLs evaluation from the MetaBrain database on “EMO-large” model.

| **Tissue** | **Option** | **ACC** | **Precision** | **Recall** | **F1-score** | **AUC** |
| --- | --- | --- | --- | --- | --- | --- |
| Hippocampus | Fine-tune | 0.568 | 0.603 | 0.837 | 0.701 | 0.579 |
|  | End-to-end | 0.457 | 0.581 | 0.367 | 0.450 | 0.415 |
| Spinal cord | Fine-tune | 0.609 | 0.577 | 0.682 | 0.625 | 0.596 |
|  | End-to-end | 0.543 | 0.563 | 0.205 | 0.300 | 0.517 |

Table S12. Single-cell eQTLs collected from the OneK1K database.

| **Cell type** | **No. of eQTLs** | **Up-regulation** | **Down-regulation** | **Middle eQTLs for training** | **Large eQTLs for training** |
| --- | --- | --- | --- | --- | --- |
| B naïve | 1,878 | 939 | 939 | 72 | 584 |
| CD4 memory | 3,544 | 1,772 | 1,772 | 74 | 386 |
| *CD4 naïve | 2,246 | 1,153 | 1,153 | 40 | 459 |
| CD8 memory | 2,306 | 1,153 | 1,153 | 98 | 396 |
| Dendritic cell | 1,956 | 978 | 978 | 28 | 293 |
| *Natural killer cell | 4,176 | 1,536 | 1,536 | 79 | 608 |

“*” indicates that the eQTLs have been balanced according to the ratio of positive and negative samples during data processing.

Table S13. The performance of EMO in predicting two disease-associated eQTL cases.

| **Disease** | **RSID** | **Gene** | **A1** | **A2** | **CHR** | **Position** | **True label** | **EMO prediction result** | **Prediction score** |
| --- | --- | --- | --- | --- | --- | --- | --- | --- | --- |
| Alzheimer’s Disease | rs1902660 | TSPAN14 | G | A | 10 | 80503927 | Up-regulation | Up-regulation | 0.937 |
| Parkinson’s Disease | rs4698412 | CD38 | G | A | 4 | 15735725 | Up-regulation | Up-regulation | 0.690 |

**Figures**

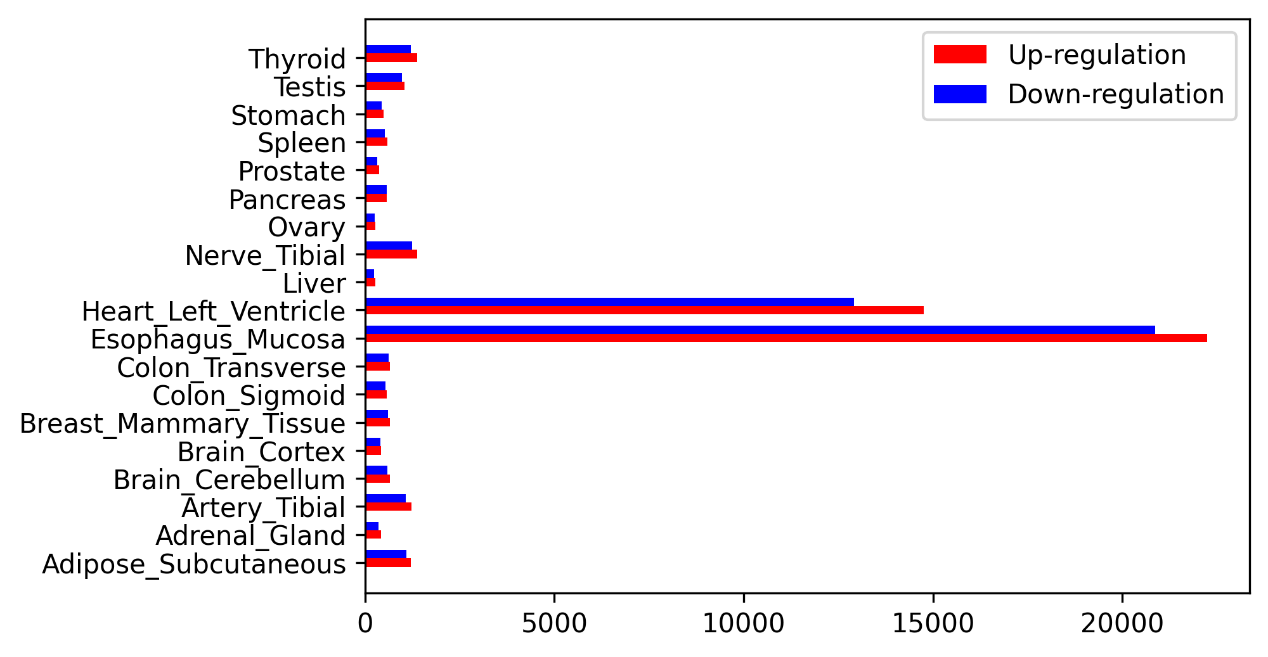

Figure S1. The distribution of positive and negative samples (‘up-regulation’ and ‘down-regulation’ eQTLs) in different tissues in the dataset before down-sampling.

\

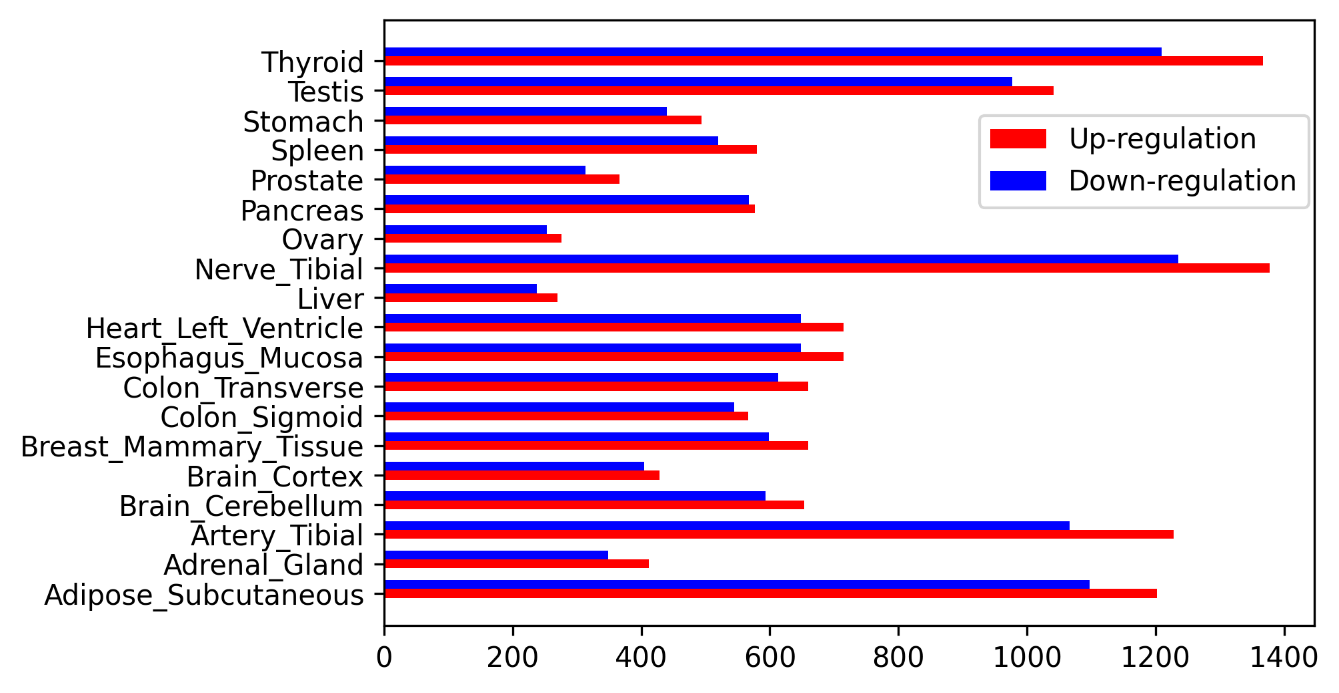

Figure S2. The distribution of positive and negative samples (‘up-regulation’ and ‘down-regulation’ eQTLs) in different tissues in the processed dataset.

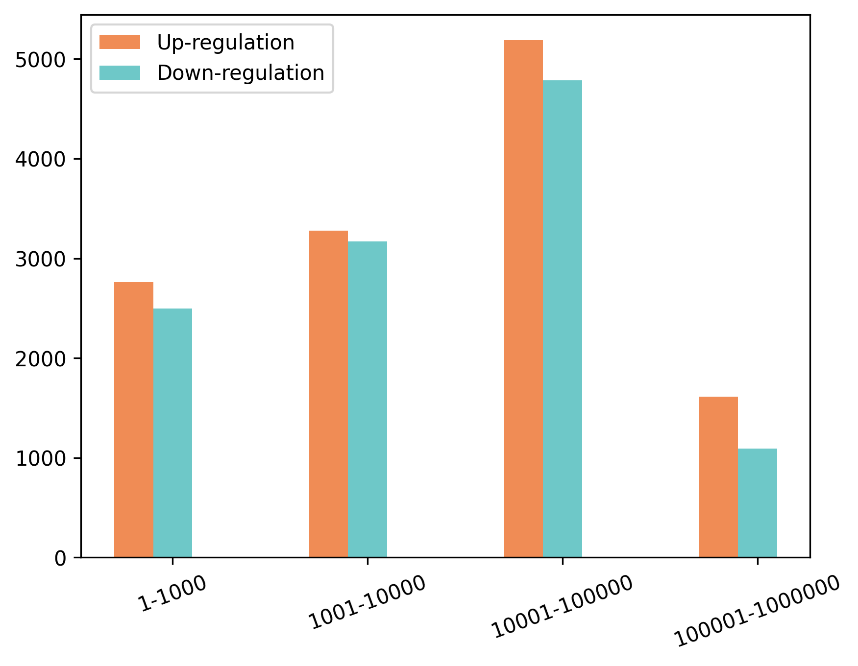

Figure S3. The distribution of different variant TSS distance of the EMO dataset.

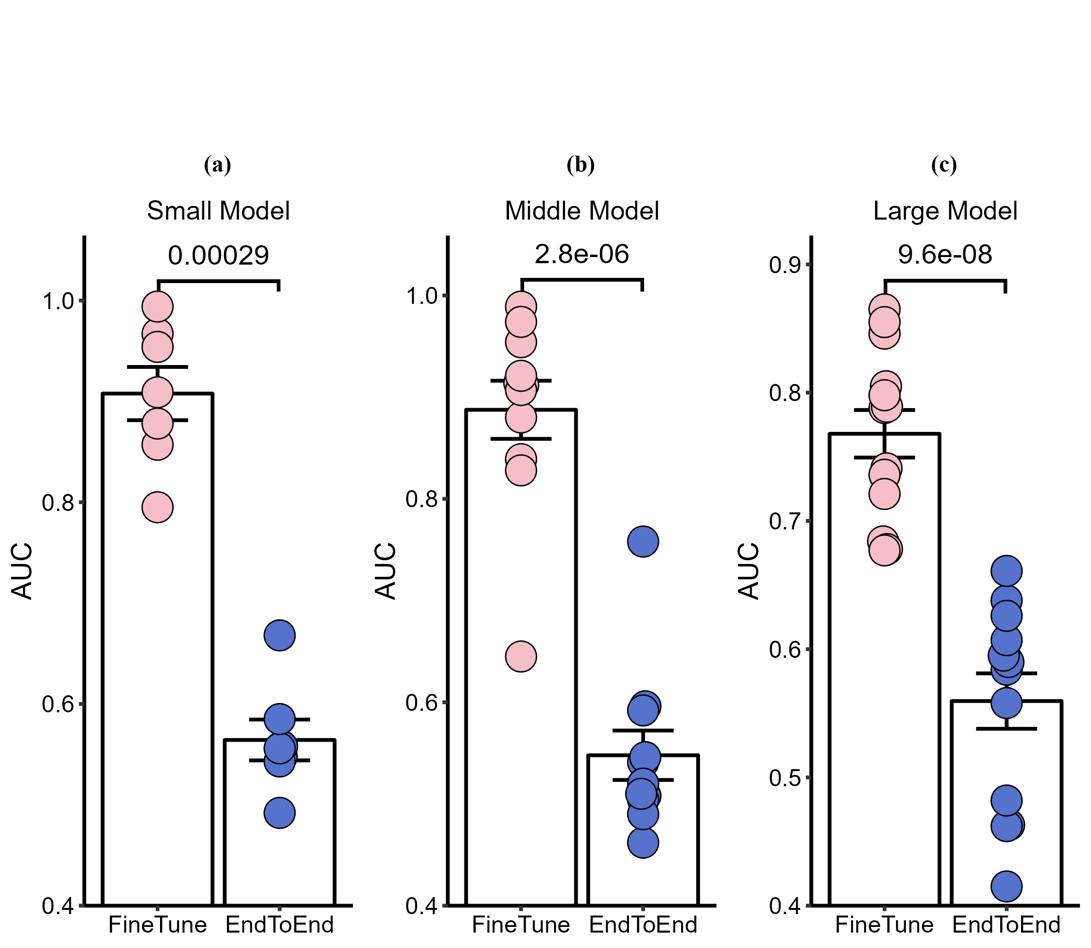

Figure S4. Tissue-specific fine-tuning and prediction for the “Small”, “Middle” and “Large” model.

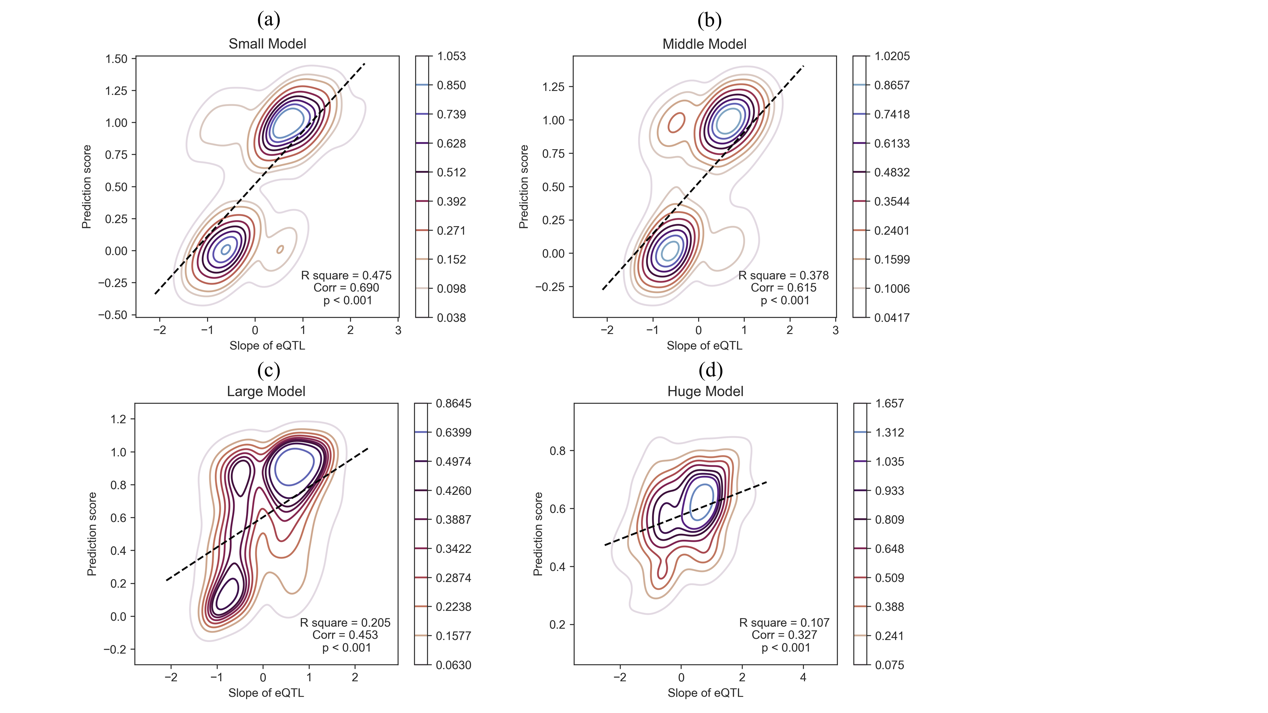

Figure S5. Slope evaluation of different EMO model, with correlation visualization between prediction score and eQTL slope.

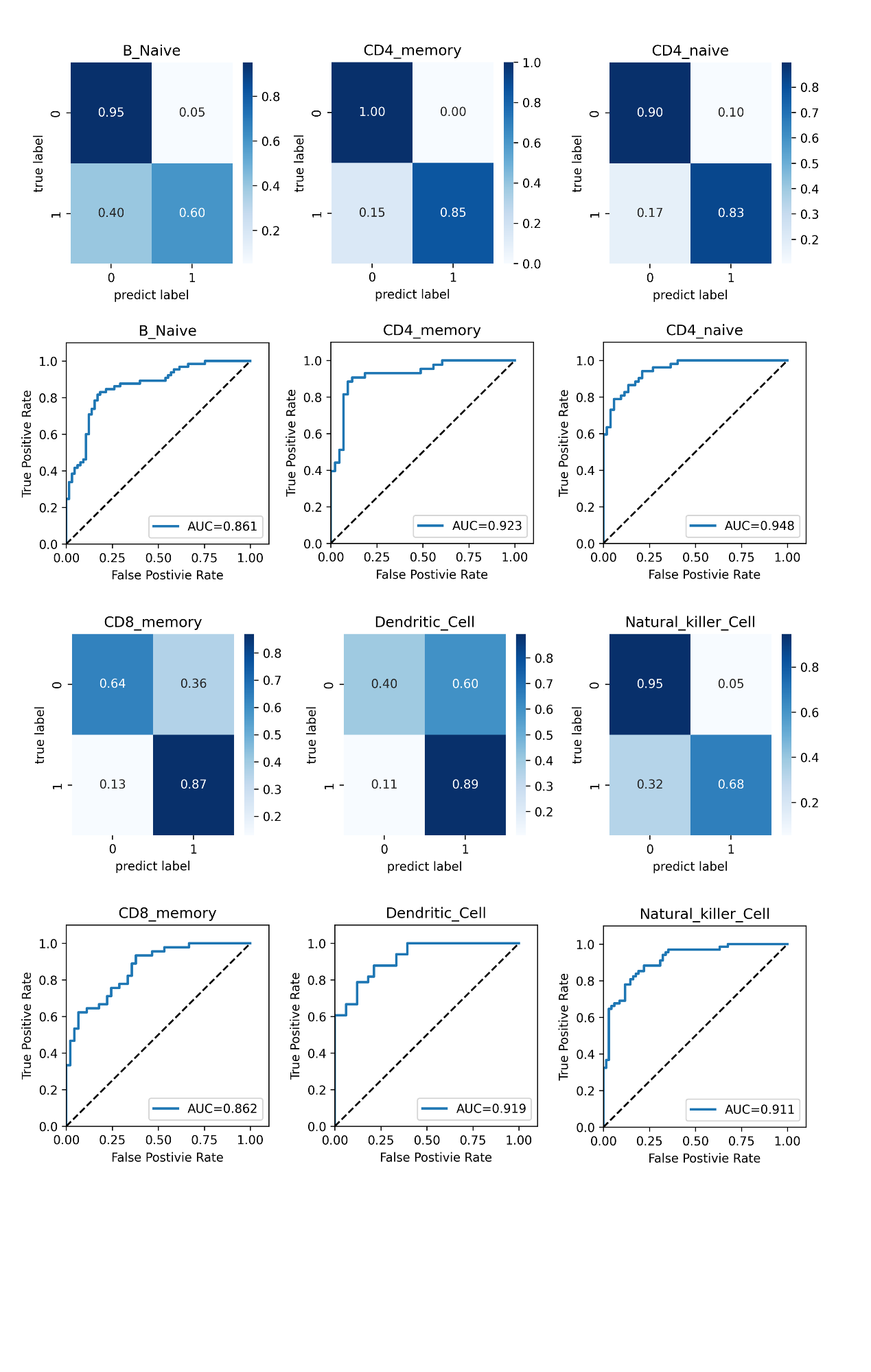

Figure S6. The prediction performance of EMO in the evaluation of six single-cell eQTLs from OneK1K.
